# Temperature and Excipient Mediated Modulation of Monoclonal Antibody Interactions Revealed by *k*D, Rheology, and Raman Spectroscopy

**DOI:** 10.1101/2025.04.06.646989

**Authors:** Olivia Noelle Eskens, Sabitoj Singh Virk, Akashdeep Singh Virk, Deepika Venkataramani, Maureen Crames, Maria Calderon Vaca, Matthew Matusewicz, Connor Smith, Joshin George, Collin Taylor, Michael S. Marlow, Samiul Amin

**Affiliations:** Department of Chemical Engineering, Manhattan University, Riverdale, NY; Department of Chemical, Environmental and Materials Engineering, University of Miami, Miami, FL; Biotherapeutics Discovery Department, Boehringer Ingelheim Pharmaceuticals Inc, Ridgefield, CT 06877

**Keywords:** Monoclonal antibody formulation, Protein-protein interactions, Diffusion interaction parameter, High-concentration viscosity, Excipient effects, Rheological characterization, Raman spectroscopy protein stability, Antibody intermolecular interactions, Subcutaneous antibody injectability, Temperature-dependent antibody interactions

## Abstract

High-concentration monoclonal antibody (mAb) formulations are frequently constrained by elevated viscosity and colloidal instability, stemming from enhanced intermolecular interactions under crowded conditions. This study delineates the thermodynamic and rheological consequences of modulating protein–protein interactions through excipient-mediated and temperature-dependent mechanisms. Using an orthogonal analytical framework comprising diffusion interaction parameter (*k*D) measurements, high-shear rheometry, and Raman spectroscopic profiling, we interrogated mAb solutions at ∼80 and 160 mg/mL across a physiologically and industrially relevant thermal window (5-45 °C). In the absence of ionic additives, high *k*D values (∼60 mL/g) indicated dominant long-range electrostatic repulsions, resulting in suppressed self-association and lower viscosity. Incorporation of NaCl (0.05% w/v) markedly decreased *k*D (∼16-20 mL/g), consistent with Debye screening of surface charges and a shift toward short-range hydrophobic and van der Waals attractions, particularly impactful at elevated protein concentrations and low temperatures. Polysorbate 20 (0.05% v/v) mitigated these interactions via preferential surface adsorption, while sucrose exhibited a dualistic, concentration-dependent influence on viscosity via preferential exclusion and entropic crowding. The combination of NaCl and PS20 yielded the most pronounced rheological suppression, reflecting synergistic attenuation of both long-range repulsion and short-range association. Raman spectral analysis of Amide I/III regions confirmed structural invariance under thermal and shear stress, attributing viscosity modulation to colloidal rather than conformational perturbations. Collectively, these data elucidate the multivariate control of interparticle potentials in mAb solutions and provide a predictive basis for engineering subcutaneous formulations that optimize manufacturability, physical stability, and injectability through strategic manipulation of colloidal interaction landscapes.

## Introduction

High-concentration subcutaneous (SC) formulations of monoclonal antibodies (mAbs) have become a linchpin of modern therapeutics, delivering large doses (>100 mg/mL) in small volumes (<1.5 mL) to facilitate patient self-administration and improve compliance^1–3^. However, as these formulations gain clinical traction, they present substantial manufacturing and delivery challenges—elevated viscosity, decreased stability, and an increased tendency toward protein aggregation—which can jeopardize both product quality and patient outcomes^4–6^. Consequently, extensive research has sought to address these hurdles, employing strategies such as optimizing formulation conditions, engineering specific excipient systems, and even altering protein sequences^7–11^.

Among these strategies, excipients play a vital role in modulating the rheological properties of mAb solutions. For instance, amino acids and sugars (*e.g.,* arginine, proline, and sucrose) can significantly affect viscosity in a concentration-dependent manner: they often increase viscosity at lower mAb concentrations yet reduce it at higher concentrations, while also stabilizing the protein’s conformation by limiting molecular dynamics^11^. Various organic, inorganic, and amino acid salts can either raise or lower solution viscosity depending on their specific interactions with mAbs^11–16^. Meanwhile, interfacial stresses, particularly at air/solid-water interfaces, can exacerbate aggregation by promoting protein adsorption and unfolding^17^. Surfactants, most notably polysorbates, help mitigate these effects by preferentially adsorbing to interfaces and, in some cases, binding directly to hydrophobic regions on the mAb^18–24^.

Temperature further complicates the formulation landscape by influencing mAb interactions and stability throughout their lifecycle, from low-temperature storage (∼5 °C) to ambient or elevated temperatures during administration^25,26^. While significantly elevated temperature excursions can induce irreversible structural changes and raise viscosity, moderate temperature increases generally lower viscosity by shifting intermolecular interactions^13,27–29^. Consequently, mapping temperature-dependent viscosity behaviors is critical for predicting and controlling mAb stability, processability, and overall formulation performance.

On a fundamental level, changes in macroscopic viscosity reflect alterations in the underlying protein-protein interaction landscape, governed by electrostatic, steric, hydrophobic, and depletion forces. Recent work underscores the importance of a delicate balance between short-range attractions and long-range repulsions (SALR) in driving mAb solution behavior under different formulation conditions^30–32^. Short-range attractions in mAb solutions are driven by hydrophobic patches, aromatic stacking, and other local interactions, while long-range repulsions are primarily due to like-charge electrostatic forces on the antibodies. In practice, this means that when two antibody molecules come very close, hydrophobic and van der Waals forces tend to pull them together, but when they are a few nanometers apart, like charges create a repulsive force that prevents them from coming too close. This combined SALR (short-range attraction, long-range repulsion) potential can explain reversible self-association (e.g., transient dimers or clusters) behavior of mAbs that typically does not result in irreversible aggregation under native conditions^33^.

In this study, we explore how temperature and excipients collectively modulate mAb interactions using an integrated analytical framework that combines *k*D measurements, rheological analysis, and Raman spectroscopy. Monoclonal antibody solutions were prepared at ∼80 mg/mL (semi-dilute condition) and 160 mg/mL (high-concentration condition), with and without specific excipients (e.g., sugars, salts, and surfactants), and their viscosity was monitored over temperatures ranging from 5 °C to 45 °C. Raman spectroscopy and *k*D analysis provided complementary insights into conformational stability and protein-protein interactions. By uniting these orthogonal methods, we illuminate the complex interplay between temperature shifts and excipient-driven effects on mAb solutions. Ultimately, our findings aim to inform the rational design of next generation, high-concentration mAb formulations that are both stable and readily manufacturable– critical objectives for advancing biotherapeutics.

## Results

### Temperature Dependence of the Diffusion Interaction Parameter (*k*D)

Measurements of the diffusion interaction parameter, *k*D, can provide insight into the balance of intermolecular forces (attractive vs. repulsive) in protein solutions, especially under varying formulation conditions. Table 1 summarizes *k*D values for the mAb solutions in 10 mM histidine buffer (pH 6) supplemented with different excipients, measured at six temperatures (5–45 °C).

**Table 1.** Diffusion interaction parameter, *k*D (mL/g), for mAb solutions containing different excipients at six temperatures.

| Temp<br>(°C) | 10 mM<br>Histidine<br>pH 6 |  | + .05% w/v<br>NaCl |  | + .05% w/v<br>Sucrose |  | + .05% v/v<br>PS20 |  | + .05% w/v<br>NaCl + .05%<br>v/v PS20 |  | + .05% w/v<br>NaCl +<br>.05% w/v<br>Sucrose |  |
| --- | --- | --- | --- | --- | --- | --- | --- | --- | --- | --- | --- | --- |
|  | Avg<br>(n=3) | SD | Avg<br>(n=3) | SD | Avg<br>(n=3) | SD | Avg<br>(n=3) | SD | Avg<br>(n=3) | SD | Avg<br>(n=3) | SD |
| 5 | 64.6 | 9.0 | 16.3 | 1.3 | 58.3 | 3.7 | 54.2 | 1.3 | 13.6 | 1.3 | 23.4 | 1.2 |
| 10 | 63.9 | 6.5 | 18.4 | 1.9 | 61.6 | 3.1 | 59.7 | 1.4 | 13.9 | 1.7 | 22.9 | 1.5 |
| 20 | 57.9 | 3.3 | 17.8 | 1.1 | 59.7 | 3.4 | 57.1 | 2.8 | 14.9 | 1.2 | 19.3 | 1.8 |
| 30 | 61.8 | 4.7 | 19.5 | 1.2 | 60.8 | 4.0 | 55.2 | 2.8 | 18.3 | 2.0 | 20.7 | 1.5 |
| 40 | 62.9 | 4.5 | 18.2 | 0.7 | 55.7 | 3.3 | 57.6 | 2.6 | 17.7 | 1.2 | 19.8 | 0.8 |
| 45 | 60.6 | 5.2 | 19.9 | 1.2 | 58.7 | 3.3 | 57.5 | 2.9 | 18.9 | 1.4 | 19.1 | 1.6 |

When the mAb was formulated in the histidine buffer alone (no added salt, sugar, or surfactant), the *k*D remained relatively high (≈58-65 mL/g), suggesting that long range repulsions were predominant. Introducing sodium chloride (NaCl (0.05% w/v)) dramatically reduced *k*D to ≈16-20 mL/g, consistent with ionic screening of electrostatic repulsions, thereby permitting stronger short-range attractive interactions. Formulations containing both NaCl and Polysorbate-20 (PS20, 0.05% v/v) had the lowest *k*D values (≈14-19 mL/g), suggesting that the combined effect of salt and surfactant shifts the balance of intermolecular forces toward a more attractive regime. One possible explanation is that NaCl screens long-range electrostatic repulsions while PS20 modifies the mAb’s surface properties, allowing short-range attractions to become relatively more influential.

In contrast, sucrose alone (0.05% w/v) did not significantly alter *k*D (≈55-61 mL/g) relative to the histidine-only control; hence, in this antibody concentration regime, sucrose did not promote short-range attractions to any large extent. However, sucrose combined with NaCl yielded intermediate *k*D values (≈19-23 mL/g), suggesting that sucrose’s preferential hydration and/or steric effects can partially offset the attractive interactions induced by salt.

Across all temperatures tested (5-45 °C), only modest shifts in *k*D were observed within each formulation, indicating that while temperature could slightly influence protein-protein interactions, the presence or absence of salt (and surfactant) remained the dominant factor dictating the net intermolecular potentials. Interestingly, all conditions exhibited a dip (local minimum) in *k*D around 20 °C, which suggested a temperature-dependent change in the balance of intermolecular forces. As the temperature increased from 10 to 20 °C, stronger short-range attractions likely developed, partially offsetting and reducing the net repulsion. By 30 °C, the balance shifted again, possibly due to increased thermal motion that restored more repulsive interactions, and hence *k*D increased. Such non-monotonic behavior, where protein interactions are most attractive at an intermediate temperature, has precedent in protein physical chemistry^34^.

Measurements of the diffusion interaction parameter (*k*D), obtained at dilute protein concentrations (1-10 mg/mL), provide insights into the nature of protein–protein interactions. High *k*D values (∼50-65 mL/g) indicate strongly repulsive interactions that effectively maintain separation between antibody molecules. Moderate decreases in *k*D (e.g., to ∼20-35 mL/g) reflect weakened yet still predominantly repulsive interactions. At semi-dilute concentrations (∼80 mg/mL), these moderate reductions typically do not result in significant viscosity increases because protein molecules remain adequately separated, limiting frequent intermolecular collisions.

However, at higher protein concentrations (≥100 mg/mL, such as 160 mg/mL), even moderate reductions in *k*D can substantially elevate solution viscosity. Under these crowded conditions, reduced long-range electrostatic repulsions allow short-range attractions, primarily hydrophobic and van der Waals interactions, to dominate, significantly increasing protein–protein encounters. Thus, formulations displaying notably lower *k*D values (e.g., those containing NaCl alone or combined with PS20) may exhibit pronounced viscosity increases at high mAb concentrations. This interpretation aligns with previous studies demonstrating that minor shifts in interaction potentials at dilute concentrations can predict substantial viscosity changes at higher protein concentrations^32,35^.

### Effect of mAb Concentration on Temperature Dependence of Viscosity

The viscosity of the mAb formulations was measured as a function of both concentration (55-160 mg/mL) and temperature (5-45 °C), as depicted in Figure 1. Generally, the solution viscosity increased with (i) increasing mAb concentration and (ii) decreasing temperature.

**Figure 1.**
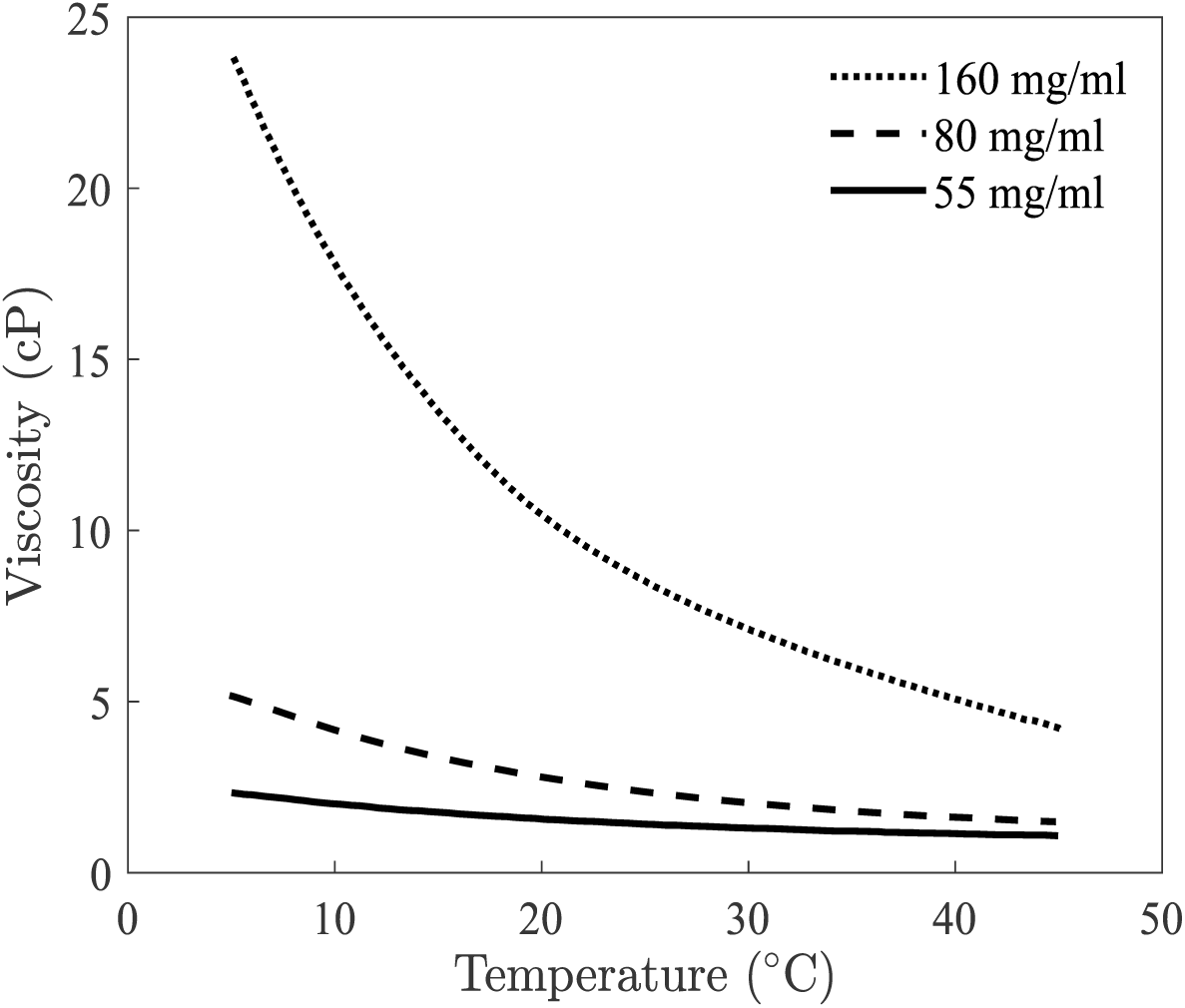
Caption: Viscosity versus temperature for various mAb concentrations (55-160 mg/mL). Error bars are smaller than the symbol size.

For dilute to semi-dilute protein solutions, the viscosity *η* may be described by a power-series expansion in terms of protein concentration c^36^:

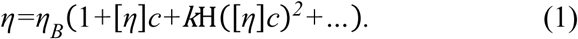

where *η_B_* is the buffer viscosity, [*η*] is the intrinsic viscosity (related to the hydrated shape and volume of the protein), and *k*H is the Huggins coefficient signifying the effect of pair interactions between proteins. At sufficiently low protein concentrations, higher-order terms become negligible, and the expression reduces to

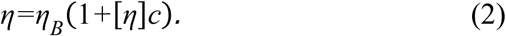

For the lowest mAb concentration studied (∼55 mg/mL), the viscosity remained nearly independent of temperature (5-45 °C) and was well-described by Eq. (2). This indicates that the shape or hydrodynamic radius of the mAb, which would affect [*η*], did not change appreciably with temperature, an inference corroborated by structural analyses (Table 2).

**Table 2.**
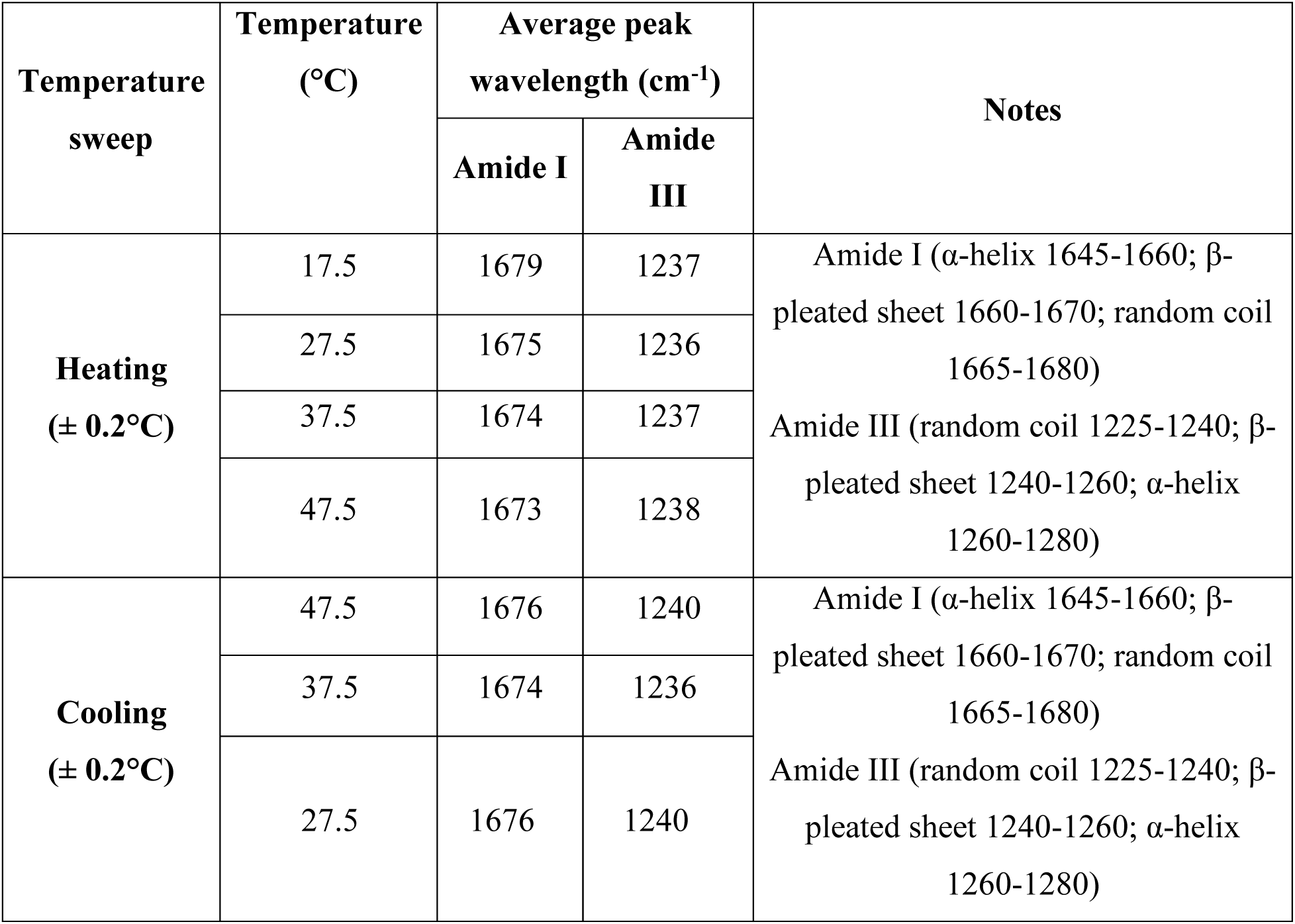
Average Raman peak positions for Amide I and Amide III regions during heating (17.5–47.5 °C) and cooling (47.5–27.5 °C).

At moderate to higher concentrations, the observed increase in viscosity, from approximately 2.4 centipoise (cP) at 80 mg/mL to about 8.5 cP at 160 mg/mL at 25 °C, reflects a significant rise in the Huggins coefficient (*k*H), signaling stronger net attractive interactions. Typically, increased temperature weakens these interactions due to enhanced thermal motion, thereby reducing viscosity^31^. However, certain protein systems may exhibit atypical behavior, with viscosity occasionally increasing at higher temperatures due to unique conformational changes, distinct aggregation pathways, or interactions with specific excipients^7^. In this study, increasing temperature consistently reduced viscosity, aligning well with the general expectation that elevated thermal energy diminishes short-range attractive interactions^31^.

### Excipients Effects on Temperature Dependence of Viscosity

Monoclonal antibodies are commonly formulated for subcutaneous injection at semi-dilute to high concentrations; excipients (*e.g.,* salts, sugars and surfactants) are often incorporated to optimize stability and reduce undesirable phenomena such as aggregation or high viscosity. In this work, the following additives were studied at a fixed level (0.05% w/v or v/v): NaCl, sucrose, and polysorbate 20 (PS20). Viscosities were measured at both semi-dilute (<100 mg/mL) and concentrated (>100 mg/mL) mAb levels ranging from 5 to 45 °C (Figure 2 and Figure 3).

**Figure 2:**
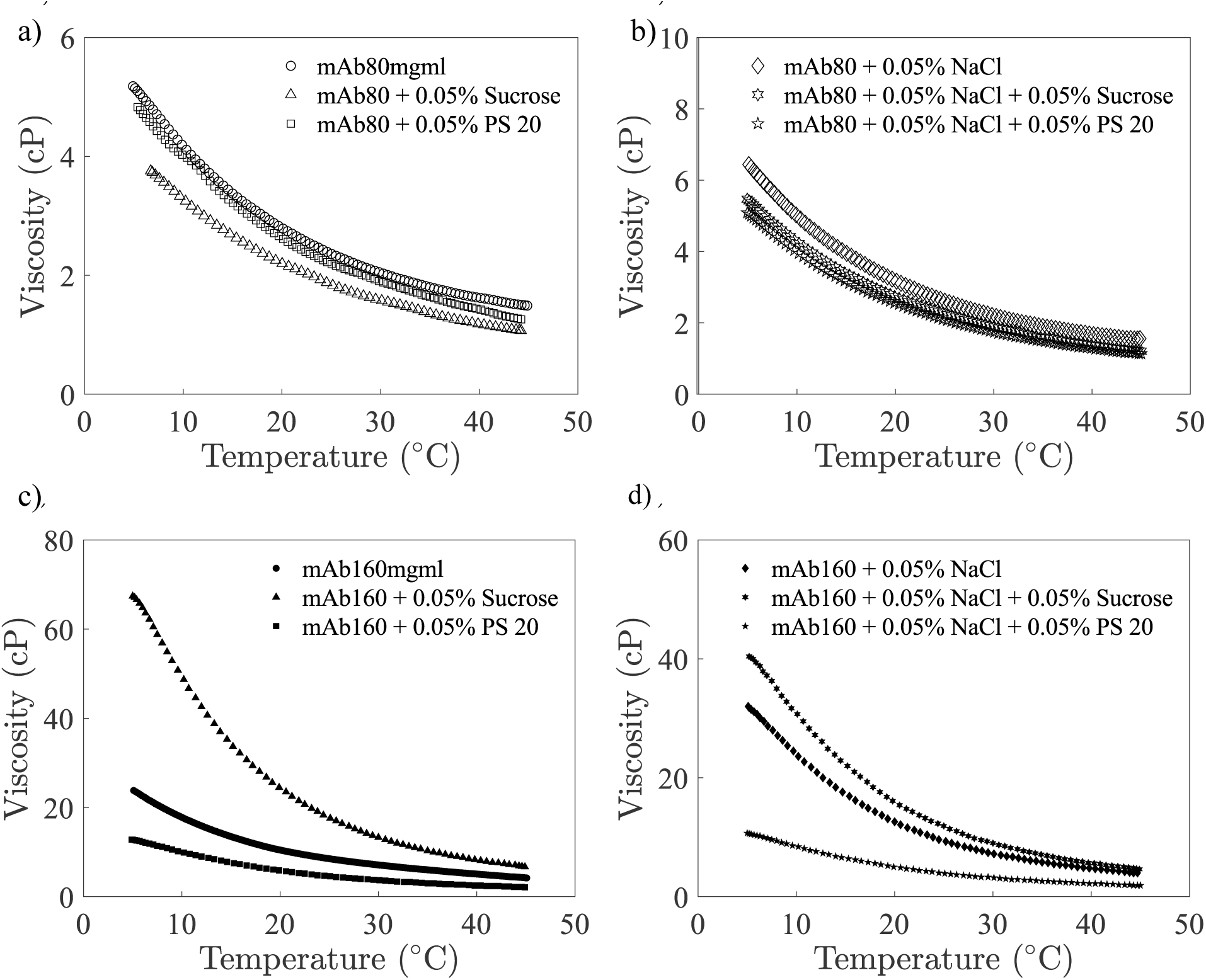
Viscosity versus temperature for mAb formulations containing excipients at semi-dilute (a,b) and concentrated (c,d) protein concentrations. Error bars are smaller than the symbol size.

**Figure 3:**
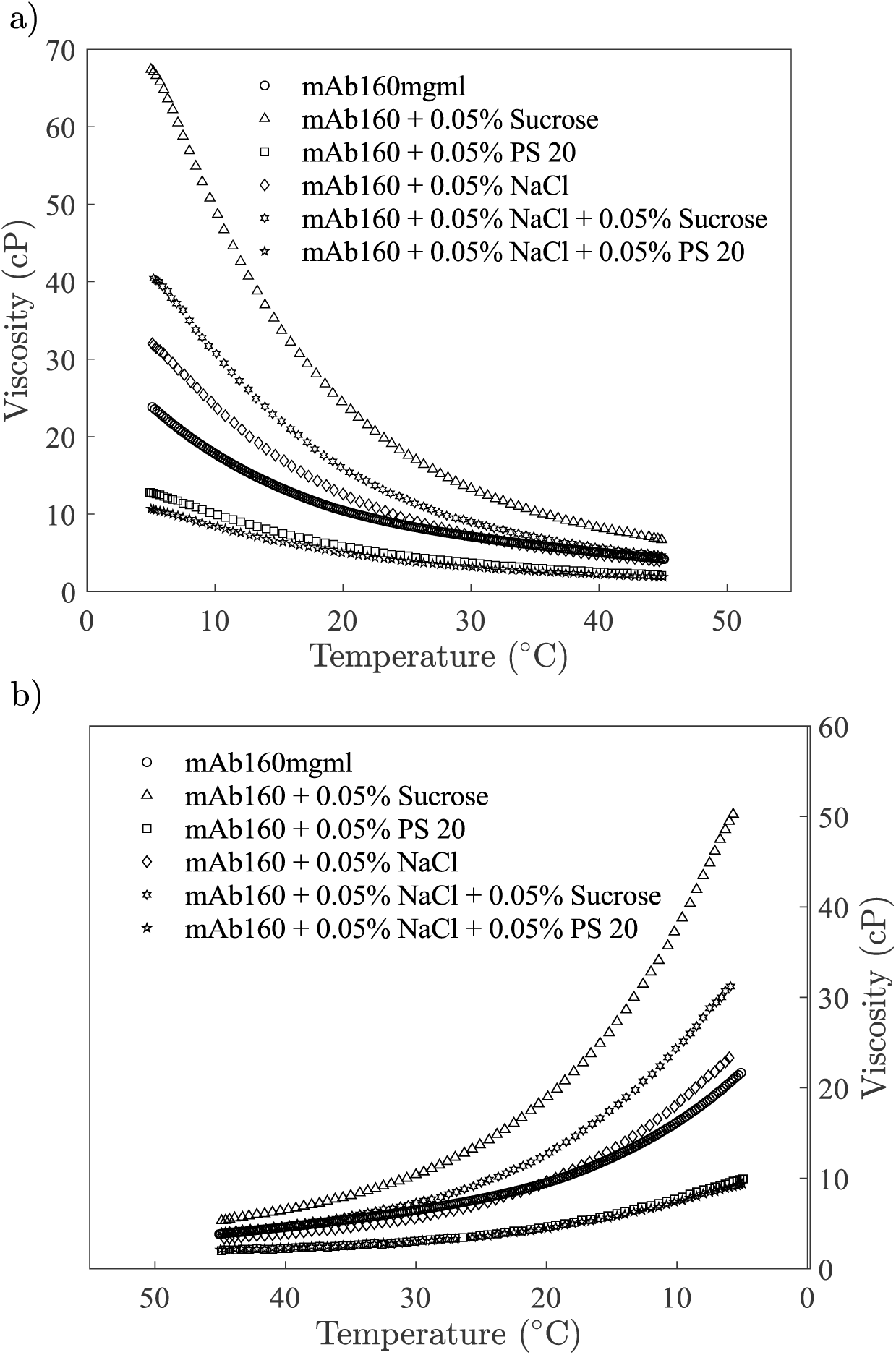
A plot of viscosity versus temperature illustrating the excipient effect on temperature dependence of viscosity during the (a) heating cycle and (b) cooling cycle. The uncertainties (one standard deviation) are smaller than the symbol size.

### Single-Component Additives

#### NaCl (0.05% w/v)

Adding NaCl to the mAb solution increased the viscosity over the entire temperature range (Figure 2b, 2d). While moderate ionic strengths can sometimes reduce viscosity,^12,35,37^ NaCl has also been shown to increase viscosity in certain protein solutions,^11,38–40^ depending on the protein’s net charge, ionic strength, and balance of electrostatic vs. hydrophobic interactions. Consistent with the *k*D results (Table 1), NaCl screened electrostatic repulsions, thereby enhancing net attractive interactions at higher mAb concentrations and/or lower temperatures.

#### Sucrose (0.05% w/v)

In semi-dilute solutions (∼80 mg/mL), sucrose addition slightly lowered viscosity compared to the pure mAb sample (Figure 2a). However, at high mAb concentration (160 mg/mL), the same sugar addition significantly increased viscosity, particularly evident at lower temperatures, highlighting a concentration-dependent effect (Figure 2c). Sucrose can stabilize proteins via preferential hydration and can also modify the effective volume or solvation shell, which may either decrease or increase viscosity depending on the mAb concentration regime^41,42^.

#### PS20 (0.05% v/v)

PS20 reduced viscosity at both semi-dilute and concentrated mAb levels, although the magnitude of reduction was more pronounced at higher concentrations (Figure 2c). Increasing mAb concentration inherently decreases the average intermolecular distance, thus diminishing electrostatic repulsion effects. For the mAb evaluated in this study, hydrophobic interactions appear to significantly contribute to intermolecular attraction as molecules become more closely packed; PS20 effectively shields these interactions. This explains why PS20 did not exhibit a similar magnitude of effect at semi-dilute and dilute concentration regimes. Nonionic surfactants such as PS20 can diminish protein–protein interactions by adsorbing to hydrophobic regions on the protein surface, thereby preventing the formation of intermolecular aggregates or films at interfaces^43–45^. However, at high shear rates (≥1,000 s⁻¹), slight increases in measured viscosity can sometimes occur if the surfactant modifies shear-thinning behavior^22^.

### Combined Excipients

Additional tests were conducted to examine formulations combining NaCl with either PS20 or sucrose:

#### NaCl + PS20

The combination of NaCl and PS20 consistently showed the lowest viscosity among all tested formulations, including those with PS20 alone (Fig. 2b and 2d). Individually, PS20 lowers viscosity by binding to antibody hydrophobic sites, preventing direct short-range hydrophobic attractions^22,46,47^. However, under conditions of low ionic strength (without NaCl), strong electrostatic repulsions significantly increase effective molecular spacing and induce structured, lattice-like ordering, elevating viscosity through electroviscous effects^37,48,49^. The introduction of moderate ionic strength via NaCl reduces these repulsions, compressing electrostatic double-layers and thus reducing structural constraints. This allows PS20 to effectively address any newly exposed short-range hydrophobic interactions, leading to the overall lowest observed viscosity^22,23,37,38^.

#### NaCl + Sucrose

In semi-dilute solutions, sucrose partially offset the viscosity increase caused by NaCl alone (Figure 2b). At high mAb concentrations, however, this same combination moderately increased viscosity relative to histidine controls, underscoring that sucrose’s effect can shift when the mAb approaches a regime of more frequent interparticle contact (Figure 2d).

Overall, raising the temperature from 5 to 45 °C decreased viscosity in all formulations, consistent with a reduction in short-range attractive interactions (as described by the SALR framework). Although viscosity decreased with increasing temperature, the magnitude of this change did not strictly mirror the modest shifts observed in *k*D, indicating that other formulation-specific factors, such as the baseline strength of protein–protein interactions and concentration effects, also play a role. Moreover, the cooling cycles (Figure 3b) did not fully retrace the heating trends (Figure 3a), suggesting a degree of hysteresis. This partial irreversibility may arise from kinetic barriers in re-establishing the original protein–protein interaction network, potentially due to subtle changes in the hydration shell or transient aggregate formation, rather than from irreversible protein unfolding or excipient precipitation^13,50,51^.

### Temperature Dependence of Secondary and Tertiary Structure of mAbs

A key goal of this study was to distinguish whether the observed viscosity changes result primarily from temperature-dependent colloidal interactions rather than large-scale conformational disruptions of the mAb. To address this, Raman spectroscopy (focusing on the Amide I and Amide III regions) was performed under the same temperature range (5-45 °C) used in the rheological experiments. Figure 4 depicts representative Raman spectra during both heating (17.5-47.5 °C) and cooling (47.5-27.5 °C).

**Figure 4:**
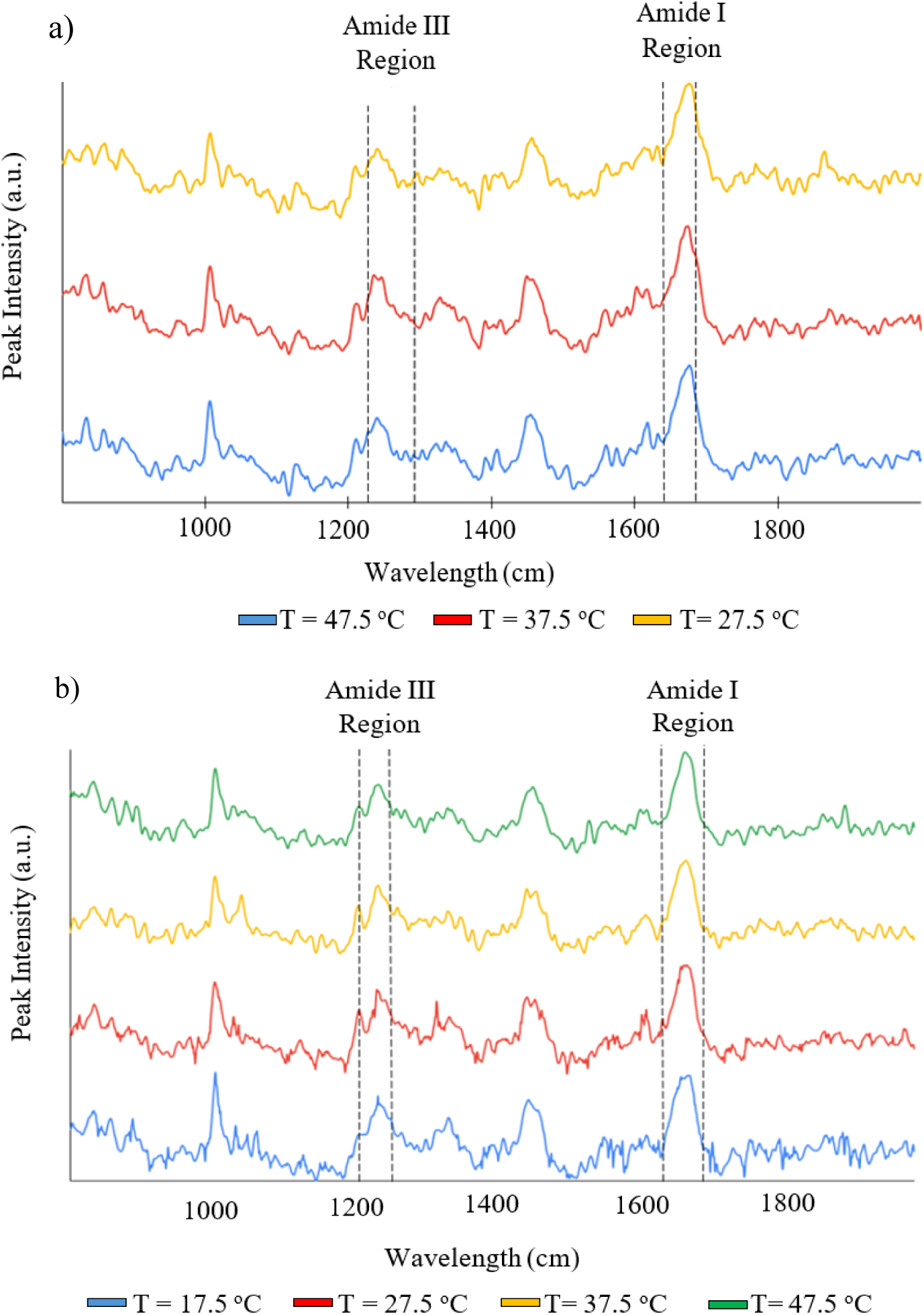
Raman spectra raw data containing secondary structure as determined by Amide I and Amide III peaks during the (a) heating cycle and (b) cooling cycle.

### Secondary Structure Analysis

Table 2 summarizes the average peak positions of the Amide I (∼1665-1680 cm^-1^ and Amide III (∼1225-1240 cm^-1^) bands. These values are characteristic of random coil or loop segments, which is consistent with immunoglobulin G domains that often adopt a mix of *β*-sheet and loop structures. Minor shifts were observed in the Amide I position (from 1679 cm^-1^ at 17.5 °C to 1673 cm^-1^ at 47.5 °C), but these were reversible upon cooling to 27.5 °C (returning to ∼1676 cm^-1^).

No significant change was noted in the Amide III region. Collectively, these data indicate that while slight rearrangements or fluctuations may occur at elevated temperature, no large-scale unfolding or transition to *β*-aggregation was detected.

### High-Shear Stability

Additional Raman measurements were conducted at high shear rates (10,000-20,000 s^-1^) to assess shear-induced conformational changes. The Amide I and Amide III bands remained essentially unchanged (Amide I at ∼1681 cm^-1^, Amide III at ∼1232-1233 cm^-1^), implying that the mAb’s secondary structure was robust under conditions well above typical manufacturing or injection shear rates. Together, these spectroscopic results confirm that the temperature-dependent viscosity changes observed in earlier sections were predominantly driven by alterations in interparticle interactions (e.g., electrostatics, hydrophobic effects) rather than significant modifications to the mAb’s secondary or tertiary structure.

## Discussion

Overall, our results highlight how the interplay between short-range attractions and long-range repulsions critically governs the colloidal and rheological behaviors of monoclonal antibody (mAb) solutions, particularly as a function of temperature, concentration, and excipient composition. By tracking the diffusion interaction parameter (*k*D) under various formulation conditions, we confirmed that ionic strength, surfactants, and osmolytes (e.g., sugars) can each modulate the net balance of intermolecular interactions that drive self-association and, consequently, influence solution viscosity.

Formulations that exhibited high *k*D values, such as the histidine control and those containing sucrose or PS20 individually, predominantly remained in a regime of long-range repulsion, which impeded close molecular contact and suppressed self-association. This is generally considered advantageous at moderate protein concentrations, where reduced self-association events typically mitigated viscosity buildup. In contrast, the pronounced decrease in *k*D upon introducing NaCl, was attributed to electrostatic screening of charges on the mAb surface. As electrostatic repulsions were weakened, even modest short-range attractions (e.g., via hydrophobic patches or Fab– Fab/Fc–Fc contacts) could drive reversible clustering, increasing solution viscosity at higher mAb concentrations.

Sucrose, while known to stabilize mAb conformations via preferential exclusion^41^, did not substantially screen electrostatic interactions and thus did not by itself induce strong short-range attractions. Nonetheless, when combined with NaCl, sucrose further lowered *k*D compared to histidine-only formulations, indicating that, once electrostatic repulsion is partially neutralized by salt, the preferential hydration effects of sugar may lead to subtle depletion forces that further shift the balance toward net attractive interactions. Similarly, PS20 alone suppressed hydrophobic interactions by binding to surface-accessible patches on the mAb, resulting in relatively higher diffusion interaction parameter (*k*D) values indicating net repulsion. However, when NaCl was added, its screening of long-range electrostatic repulsions allowed antibodies to approach more closely, thereby revealing residual hydrophobic attractions that remained even with PS20 present. This synergistic effect led to a markedly lower *k*D in formulations containing both NaCl and PS20 compared to those with PS20 alone^38,52–54^. Such multifactorial interplay underscores the necessity of judiciously optimizing ionic strength, surfactant levels, and sugar content to achieve the desired colloidal stability.

The concentration dependence of mAb solution viscosity became evident when examining dilute (∼ 55 mg/mL) versus higher-concentration regimes under varying temperature. In dilute solutions, intermolecular interactions were minimal due to the large average distance between mAb molecules, and the solution viscosity primarily reflected the intrinsic properties of the mAb (e.g., shape and hydrodynamic radius), which remained largely invariant with temperature in the range studied.

As the mAb concentration approached semi-dilute and concentrated regimes, pairwise and higher-order interactions became significant, leading to an increase in the Huggins coefficient (*k*H). Analytical approaches based on short-range attractive and long-range repulsive (SALR) interactions provide a framework to compute *k*H, and by extension, the solution viscosity (η), using parameters such as the inverse Debye length (*κ*), short-range attraction strength (*1/τ\**), and long-range repulsion strength (*α*)^55^. In systems near the isoelectric point or under partial electrostatic screening, short-range attractions could dominate and dramatically increase *k*H (and thus viscosity), as reported in previous studies^32,56^. Our findings corroborated this concept, particularly under conditions that favored hydrophobic driven clustering.

Adding NaCl to the mAb solution reinforced the notion of SALR-driven viscosity. Low to moderate salt concentrations screened long-range repulsions more effectively than short-range attractions, thereby unmasking or amplifying attractive interactions that raised viscosity^32^. In our experiments, even 0.05% (w/v), or 8.6 mM, NaCl increased viscosity confirming that electrostatic screening triggered short-range attractions to become the dominant driver of rheological behavior. The effect of sucrose on solution viscosity depended strongly on the protein concentration. At low concentrations, sucrose was preferentially excluded from the protein surface, reducing hydrophobic contacts and lowering viscosity. However, at higher mAb concentrations, the additional osmolyte could effectively crowd protein molecules, enhancing intermolecular contacts and increasing viscosity, even under conditions where sucrose had been beneficial at lower concentrations. This dual behavior aligned with earlier observations that sugars could strengthen hydrophobic interactions (i.e., short-range attractions) under crowded conditions^42^, thereby amplifying or diminishing viscosity depending on the nature and density of protein packing (Figure 2). As described in the results section, PS20 alone effectively mitigated hydrophobic attractions by binding to exposed/surface-accessible mAb regions, thereby lowering viscosity. In formulations with added salt, the screening of long-range electrostatic repulsions further compressed the effective molecular spacing, enabling PS20 to interact more efficiently with hydrophobic sites. Consequently, the combination of NaCl and PS20 yielded the lowest viscosity among all tested formulations (Fig. 2b and 2d), as it reduced both the repulsive and the residual attractive forces between antibody molecules. Such synergistic or antagonistic outcomes between surfactants, salts, and sugars illustrated the complex landscape of excipient optimization for mAb formulations.

In all excipient-containing systems, temperature shifts altered the balance between short-range attractions and long-range repulsions, further modulating viscosity. Although our data suggested that no major structural rearrangements occurred (see Raman discussion below), thermal perturbations to local protein–protein interaction energies could still be sufficient to influence viscosity^57^.

Raman spectroscopy corroborated that the mAb retained a predominantly random coil–like secondary structure across a broad temperature range (∼4-40 °C) and under high shear conditions relevant to manufacturing (e.g., pumping, filtration) and subcutaneous injection. The minimal shifts in Amide I (1673-1681 cm⁻¹) and Amide III (1232-1240 cm⁻¹) regions remained well within published ranges for random coil signatures, indicating minimal conformational changes. Consequently, any viscosity fluctuations were more plausibly governed by intermolecular colloidal forces rather than by large-scale unfolding or tertiary structural transitions. This structural resilience in mAb solutions offers important practical benefits: *Manufacturing Flexibility*: Processes involving high shear (pumping, mixing, filtration) are less likely to induce denaturation or irreversible aggregation. *Clinical Delivery*: Subcutaneous injection at high protein concentrations is feasible without concern for shear-induced conformational disruptions. *Thermal Robustness*: The absence of major thermal transitions simplifies storage and shipping considerations, reducing the risk of viscosity spikes or irreversible aggregation driven by unfolding events. Moreover, knowing that the mAb backbone remains largely unaltered by these stresses allows formulation scientists to focus on fine-tuning ionic strength, surfactant concentration, and sugar content to control viscosity, without having to mitigate unfolding-related aggregation pathways.

From a manufacturing and delivery standpoint, especially for high-concentration subcutaneous injections, excessively low *k*D (and by extension, strong net attraction) poses a clear risk for elevated viscosity. The rising prevalence of self-administration and pen-injector systems further underscores the need to maintain mAb formulations at a manageable viscosity level. Our findings emphasize that balancing ionic strength and carefully selecting excipients (e.g., surfactants, sugars) is crucial for sustaining sufficient long-range repulsive forces to keep viscosity within acceptable bounds for both manufacturing operations and patient delivery. These findings may be relevant to other antibodies with similar biophysical characteristics, though antibody-specific differences are significant^56^. The trends observed here, e.g. salt weakening repulsion or sugar-induced crowding, are consistent with general protein interaction principles, but the exact magnitude and balance of forces can vary from one mAb to another^59^.

Ultimately, an integrated understanding of how *k*D, *k*H, and solution viscosity respond to temperature and excipient combinations can guide the design of robust, stable, and clinically practical mAb formulations. Formulators can leverage these insights to strategically manipulate colloidal interactions, using salts, sugars, and surfactants, to achieve the dual goals of maintaining protein stability and controlling viscosity under real-world storage, handling, and administration conditions. Future work could examine additional excipients. For instance, amino acids like arginine or lysine have been shown to disrupt attractive mAb-mAb interactions and significantly lower solution viscosity^60^. Conversely, incorporating inert crowding agents (e.g., small polyethylene glycol) could be used to deliberately enhance depletion attractions and test the robustness of the formulation^58^. Exploring such excipients would further elucidate the balance of forces in mAb solutions and could identify new strategies to control viscosity and stability.

In summary, this work demonstrates how short-range attractions and long-range repulsions compete to shape mAb solution viscosity, with ionic strength, sugar, and surfactant additives each exerting distinct and sometimes synergistic effects on colloidal stability. Temperature variations further modulate these interactions, although the mAb retains its higher-order structural integrity. By dissecting the drivers of *k*D and *k*H and correlating them with viscosity data, we provide a roadmap for tuning mAb formulations to balance stability, manufacturability, and injectability— key goals in the development of next-generation high-concentration biotherapeutics.

Below is a table synthesizing the key points of our study into practical formulation guidelines:

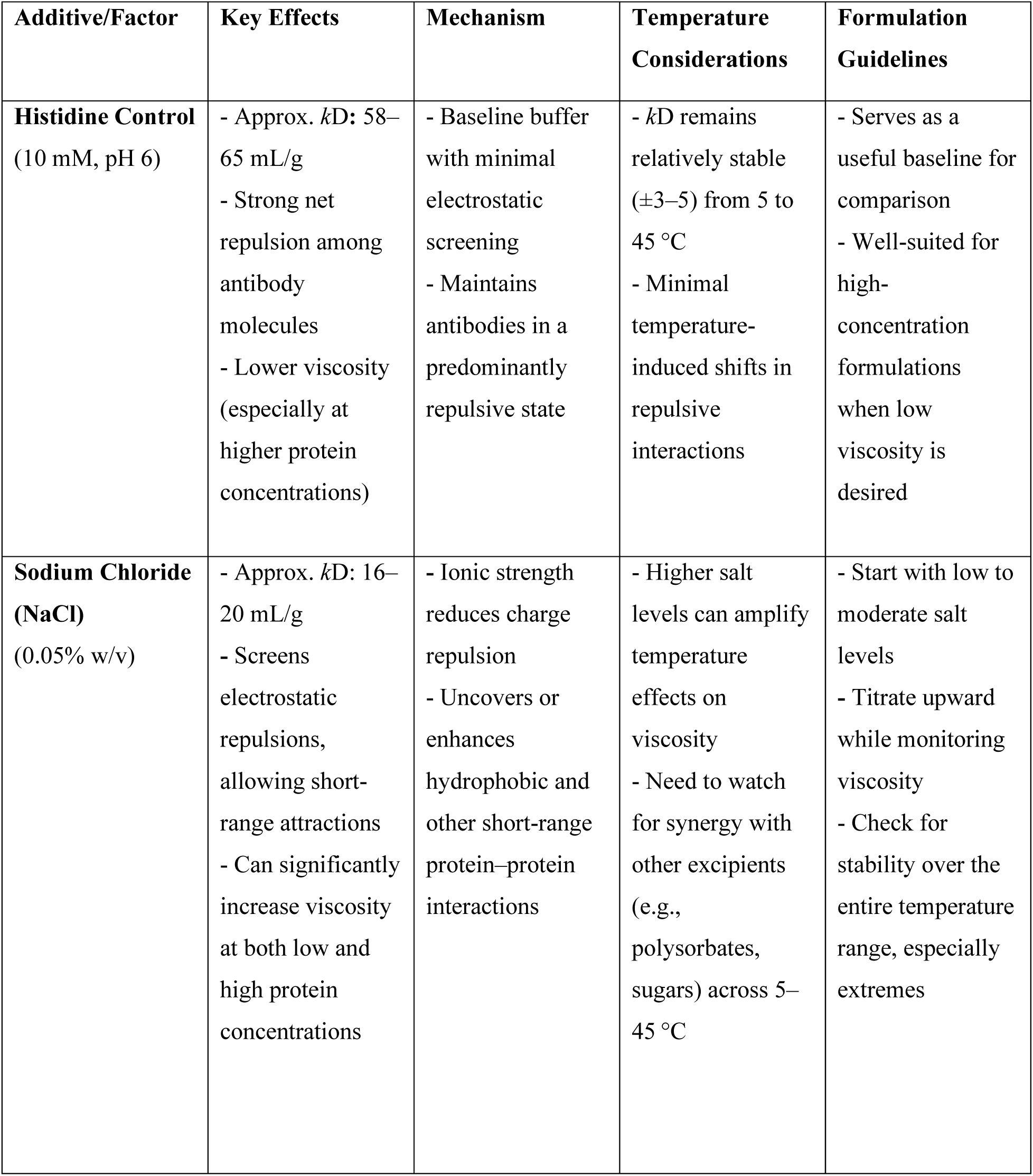
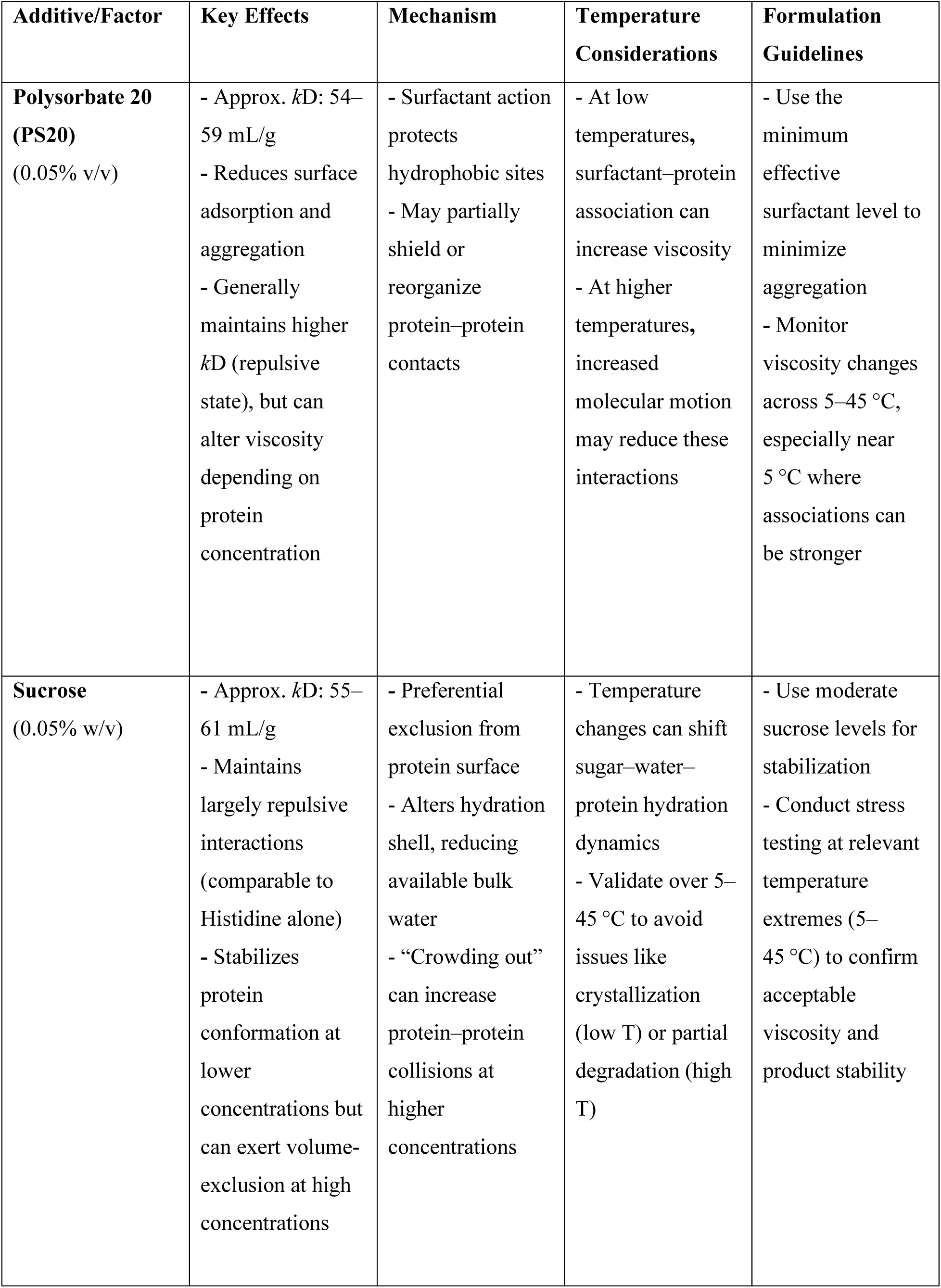
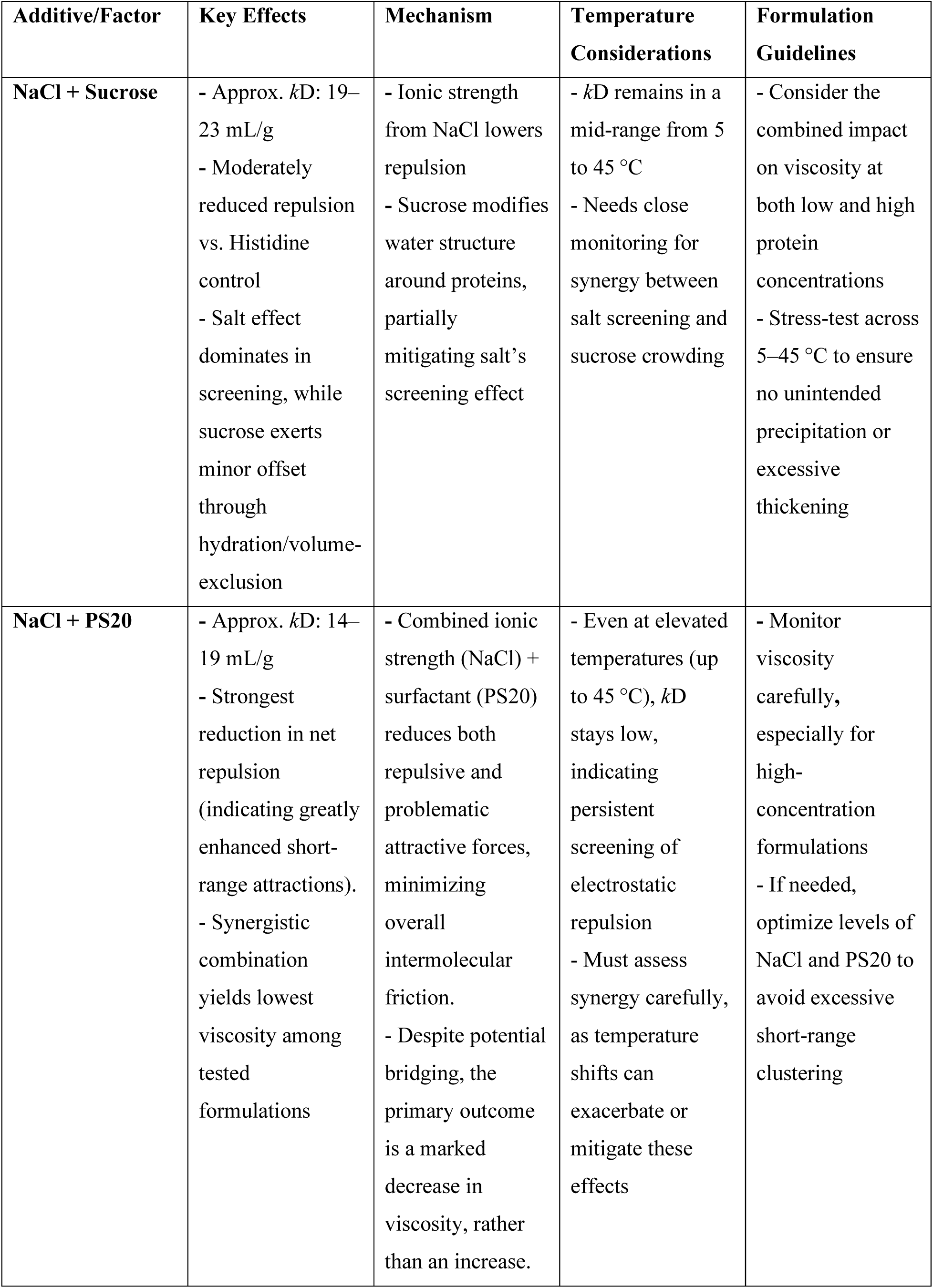
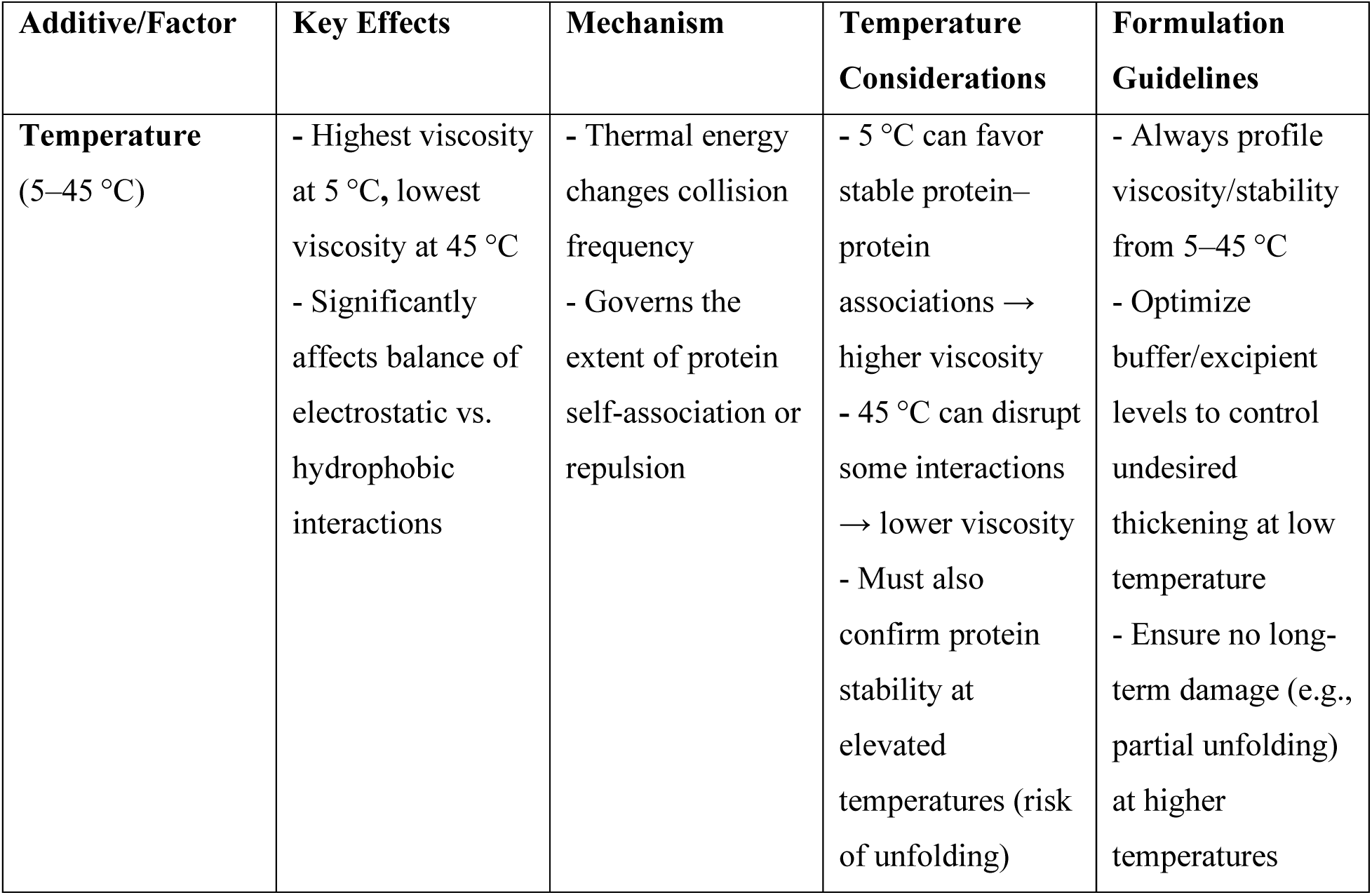

**Possible use of the table:**

1. **Identifying the Baseline**: Histidine control serves as a reference for high *k*D (strong net repulsion) and relatively low viscosity.
2. **Evaluating Salt Requirements**: NaCl or combined ‘NaCl + excipients’ can drastically lower *k*D, screening electrostatic repulsion and boosting short-range attractions. This may be acceptable at low protein concentrations but risky at high concentrations (viscosity spikes).
3. **Considering Carbohydrate Stabilizers**: Sucrose maintains a higher *k*D alone, but combined with NaCl can still lower net repulsion. Validate thoroughly at both low and high protein loads.
4. **Surfactant Choices**: PS20 effectively reduces aggregation but can sometimes contribute to unexpected viscosity changes, especially in salted solutions or at low temperatures.
5. **Temperature Profiling**: Across 5-45 °C, always verify that the formulation neither becomes too viscous at low temperatures nor unstable at high temperatures. Each additive can behave differently under thermal stress.
6. **Iterate and Refine:** Combine analytical data (particle counts, turbidity, interaction parameter, rheology) to fine-tune the buffer and excipient composition, ensuring acceptable viscosity and protein stability across the intended temperature range.

## Materials and Methods

### Materials

The monoclonal antibody utilized in this study was supplied by Boehringer Ingelheim at a concentration of 160 mg/mL in a histidine buffer solution stabilized at a pH of 6. In this investigation, three excipients were incorporated: (i) sucrose, (ii) NaCl, and (iii) polysorbate 20. Both sucrose and NaCl were procured from Fischer Chemicals and were formulated into the mAb solution at a concentration of 0.05% (w/v). Meanwhile, the surfactant polysorbate 20, provided by Croda Inc., was initially available as a stock solution at a concentration of 0.05% (v/v).

### Sample preparation

For this study, all samples were prepared and measured immediately after preparation. The sample preparation method varied depending on the specific property under investigation. For viscosity measurements in the presence of excipients, mAb samples with an initial concentration of 160 mg/mL (provided by Boehringer Ingelheim) were diluted using a histidine buffer solution to achieve a final concentration of 80 mg/ml whenever required. In the case of diffusion interaction parameter (*k*D) measurements, both protein solutions with and without excipients were diluted to concentrations ranging between 1 and 10 mg/mL; this was accomplished by employing the histidine buffer solution for dilution. For Raman spectroscopy measurements, a 55 mg/mL mAb sample was utilized, which was prepared by diluting the 160 mg/mL stock solution with the same histidine buffer solution.

### Viscosity Measurements

The Discovery HR-3 mechanical rheometer (TA Instruments, Delaware, USA) was employed to measure the viscosity of the protein formulations. In this approach, a 40 mm parallel plate geometry was utilized for all rheological measurements. The plates were brought into contact with the sample at an 80 μm gap, as determined by a prior gap dependency study^61^. For the present work, this narrow gap was deliberately chosen to achieve high shear rates and to alleviate artifacts arising from protein structuring at the air-water interface, as demonstrated by Hirschman et al.^61^. Viscosity measurements were conducted over a temperature range of 5 °C to 45 °C at a constant shear rate of 1000 s⁻¹. This process was subsequently reversed from 45 °C down to 5 °C, and then repeated once more from 5 °C up to 45 °C. After the desired gap and shear rate were established, both the parallel plate geometry and the sample were covered with circular metal plates to prevent drying and potential solvent loss, thereby serving as an effective barrier^62^.

Shear rate, defined as the ratio of flow velocity to gap distance, can be increased either by augmenting the flow velocity within the geometry or by reducing the gap distance^61–63^. To ensure reproducibility and reliability, all measurements were repeated at least three times. The sample volume required for each trial was calculated based on the geometry size and the set gap. However, it has been shown that employing small sample sizes and narrow gaps can introduce potential gap errors. Such errors may result from possible misalignment or other artifacts that could influence the accuracy of the results. To account for this potential error, an additional 50 μL of sample was added when using small gaps, intentionally creating a slight overfill^61,64,65^.

### Diffusion Interaction Parameter Methodology

The diffusion interaction parameter, *k*D was measured using a dynamic light scattering technique. Specifically, a Malvern Zetasizer Ultra (Malvern, U.K.) was employed to elucidate the relationship between the diffusion coefficient, *D_m_*, and the sample concentration, *c*, as described by equation (3), where *D*₀ represents the infinite-dilution diffusion coefficient^66^.

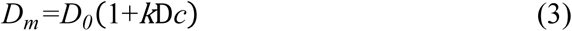

For the experiment, protein concentrations ranged between 1 and 10 mg/mL. A linear regression analysis between concentration and *D_m_* was performed at six different temperatures (5, 10, 20, 30, 40, and 45 °C). These six measurements were strategically selected to acquire diffusion coefficient data corresponding to the temperature ramp employed in the viscosity measurement study. Both the y-intercept and the slope of the linear regression were utilized to determine the value of *k*D. The *D_m_* values were computed using the ZS XPLORER software. A disposable plastic cuvette, containing 1.2 mL of the diluted sample, was used for each measurement. After a 5-minute equilibration period at the specified temperature, 30 size measurements were recorded per sample, with each run lasting 3.36 seconds. The average diffusion coefficient was then determined for each sample and used in the computation of *k*D.

### Raman Spectroscopy Measurements

The Discovery HR-3 TA Instruments (Delaware, USA) mechanical rheometer—equipped with a Rheo-Raman accessory and an integrated iXR Raman Spectrometer from Thermo Fisher Scientific, was employed to thoroughly investigate the secondary and tertiary structural changes in monoclonal antibody (mAb) samples. This sophisticated setup featured a 532 nm laser calibrated at 10 mW, and the measurements were conducted using a 55 mg/mL antibody solution prepared by diluting a 160 mg/mL stock mAb solution with a histidine buffer. In this work, a detailed analysis of secondary structural modifications was developed by subjecting the samples to both isobaric and isochoric heating and cooling cycles, as well as to high shear conditions.

Secondary structural thermal stability experiments were performed under atmospheric pressure isobaric conditions using a constant volume of approximately 1 mL of solution. The plate temperature was increased in 5 °C increments from 17.5 °C to 52.5 °C, with each interval maintained for roughly 15 minutes to ensure that the sample reached thermal equilibrium. During each temperature hold, spectra were recorded every 5 minutes. Sample conditions were carefully verified throughout the experiments to confirm that the solution remained in a liquid state and did not undergo any phase transformation or film formation. An analogous procedure was carried out for the cooling experiments, with the temperature decreasing from 52.5 °C to 18.1 °C; although the target end temperature of 17.5 °C was not achieved due to equipment and environmental constraints. Additionally, shear rates of 10,000 s⁻¹ and 20,000 s⁻¹ were selected to mirror the high shear conditions observed during typical antibody injections and to evaluate protein stability under the rapid mixing conditions encountered in industrial processes. During the high shear experiments, exposures were recorded at 10-minute intervals over a period of 30 minutes.

## Acknowledgements

In memoriam of a dear colleague, Olivia Noelle Eskens, who made important and significant contribution to this work.

## Disclosure statement

The authors report there are no competing interests to declare.

## Funding

This research was funded by Boehringer Ingelheim.

## Notes

### Competing Interest Statement

The authors have declared no competing interest.

## References

1. Harris RJ, Shire SJ, Winter C. Commercial manufacturing scale formulation and analytical characterization of therapeutic recombinant antibodies. Drug development research. 2004 Mar;61(3):137–54.

2. Shire, SJ. 6 – Challenges in the subcutaneous (SC) administration of monoclonal antibodies (mAbs), in Monoclonal Antibodies. Woodhead Publishing. 2015 Apr 24;131–138.

3. Yadav S, Shire SJ, Kalonia DS. Factors affecting the viscosity in high concentration solutions of different monoclonal antibodies. Journal of pharmaceutical sciences. 2010 Dec 1;99(12):4812–29.

4. Saluja A, Kalonia DS. Nature and consequences of protein–protein interactions in high protein concentration solutions. International journal of pharmaceutics. 2008 Jun 24;358(1-2):1–5.

5. Shire SJ, Shahrokh Z, Liu JU. Challenges in the development of high protein concentration formulations. Journal of pharmaceutical sciences. 2004 Jun 1;93(6):1390–402.

6. Daugherty AL, Mrsny RJ. Formulation and delivery issues for monoclonal antibody therapeutics. Advanced drug delivery reviews. 2006 Aug 7;58(5-6):686–706.

7. Dear BJ, Hung JJ, Truskett TM, Johnston KP. Contrasting the influence of cationic amino acids on the viscosity and stability of a highly concentrated monoclonal antibody. Pharmaceutical research. 2017 Jan;34:193–207.

8. Hung JJ, Dear BJ, Dinin AK, Borwankar AU, Mehta SK, Truskett TT, Johnston KP. Improving viscosity and stability of a highly concentrated monoclonal antibody solution with concentrated proline. Pharmaceutical Research. 2018 Jul;35:1–4.

9. Yadav S, Sreedhara A, Kanai S, Liu J, Lien S, Lowman H, Kalonia DS, Shire SJ. Establishing a link between amino acid sequences and self-associating and viscoelastic behavior of two closely related monoclonal antibodies. Pharmaceutical research. 2011 Jul;28:1750–64.

10. Yadav S, Shire SJ, Kalonia DS. Factors affecting the viscosity in high concentration solutions of different monoclonal antibodies. Journal of pharmaceutical sciences. 2010 Dec 1;99(12):4812–29.

11. Binabaji E, Ma J, Zydney AL. Intermolecular interactions and the viscosity of highly concentrated monoclonal antibody solutions. Pharmaceutical research. 2015 Sep;32:3102–9.

12. Kanai S, Liu JU, Patapoff TW, Shire SJ. Reversible self-association of a concentrated monoclonal antibody solution mediated by Fab–Fab interaction that impacts solution viscosity. Journal of pharmaceutical sciences. 2008 Oct 1;97(10):4219–27.

13. Godfrin PD, Zarraga IE, Zarzar J, Porcar L, Falus P, Wagner NJ, Liu Y. Effect of hierarchical cluster formation on the viscosity of concentrated monoclonal antibody formulations studied by neutron scattering. The Journal of Physical Chemistry B. 2016 Jan 21;120(2):278–91.

14. Du W, Klibanov AM. Hydrophobic salts markedly diminish viscosity of concentrated protein solutions. Biotechnology and bioengineering. 2011 Mar;108(3):632–6.

15. Larson AM, Weight AK, Love K, Bonificio A, Wescott CR, Klibanov AM. Bulky polar additives that greatly reduce the viscosity of concentrated solutions of therapeutic monoclonal antibodies. Journal of pharmaceutical sciences. 2017 May 1;106(5):1211–7.

16. Wang S, Zhang N, Hu T, Dai W, Feng X, Zhang X, Qian F. Viscosity-lowering effect of amino acids and salts on highly concentrated solutions of two IgG1 monoclonal antibodies. Molecular pharmaceutics. 2015 Dec 7;12(12):4478–87.

17. Shieh IC, Patel AR. Predicting the agitation-induced aggregation of monoclonal antibodies using surface tensiometry. Molecular pharmaceutics. 2015 Sep 8;12(9):3184–93.

18. Bos MA, Van Vliet T. Interfacial rheological properties of adsorbed protein layers and surfactants: a review. Advances in colloid and interface science. 2001 Jul 27;91(3):437–71.

19. Khan TA, Mahler HC, Kishore RS. Key interactions of surfactants in therapeutic protein formulations: a review. European journal of pharmaceutics and biopharmaceutics. 2015 Nov 1;97:60–7.

20. Rippner Blomqvist B, Ridout MJ, Mackie AR, Wärnheim T, Claesson PM, Wilde P. Disruption of Viscoelastic β-Lactoglobulin Surface Layers at the Air− Water Interface by Nonionic Polymeric Surfactants. Langmuir. 2004 Nov 9;20(23):10150–8.

21. Kapp SJ, Larsson I, Van De Weert M, Cárdenas M, Jorgensen L. Competitive adsorption of monoclonal antibodies and nonionic surfactants at solid hydrophobic surfaces. Journal of pharmaceutical sciences. 2015 Feb 1;104(2):593–601.

22. Patapoff TW, Esue O. Polysorbate 20 prevents the precipitation of a monoclonal antibody during shear. Pharmaceutical development and technology. 2009 Dec 1;14(6):659–64.

23. Wang S, Wu G, Zhang X, Tian Z, Zhang N, Hu T, Dai W, Qian F. Stabilizing two IgG1 monoclonal antibodies by surfactants: balance between aggregation prevention and structure perturbation. European Journal of Pharmaceutics and Biopharmaceutics. 2017 May 1;114:263–77.

24. Tein YS, Zhang Z, Wagner NJ. Competitive surface activity of monoclonal antibodies and nonionic surfactants at the air–water interface determined by interfacial rheology and neutron reflectometry. Langmuir. 2020 Jun 17;36(27):7814–23.

25. Shire SJ. Formulation and manufacturability of biologics. Current opinion in biotechnology. 2009 Dec 1;20(6):708–14.

26. Yang Y, Velayudhan A, Thornhill NF, Farid SS. Multi-criteria manufacturability indices for ranking high-concentration monoclonal antibody formulations. Biotechnology and Bioengineering. 2017 Sep;114(9):2043–56.

27. Tang Q, Munro PA, McCarthy OJ. Rheology of whey protein concentrate solutions as a function of concentration, temperature, pH and salt concentration. Journal of Dairy Research. 1993 Aug;60(3):349–61.

28. Josephson LL, Galush WJ, Furst EM. Parallel temperature-dependent microrheological measurements in a microfluidic chip. Biomicrofluidics. 2016 Jul 14;10(4):043503.

29. Fukuda M, Watanabe A, Hayasaka A, Muraoka M, Hori Y, Yamazaki T, Imaeda Y, Koga A. Small-scale screening method for low-viscosity antibody solutions using small-angle X-ray scattering. European Journal of Pharmaceutics and Biopharmaceutics. 2017 Mar 1;112:132–7.

30. Hong T, Iwashita K, Shiraki K. Viscosity control of protein solution by small solutes: a review. Current Protein and Peptide Science. 2018 Aug 1;19(8):746–58.

31. Woldeyes MA, Qi W, Razinkov VI, Furst EM, Roberts CJ. Temperature Dependence of Protein Solution Viscosity and Protein–Protein Interactions: Insights into the Origins of High-Viscosity Protein Solutions. Molecular Pharmaceutics. 2020 Nov 10;17(12):4473–82.

32. Virk SS, Underhill PT. Application of a Simple Short-Range Attraction and Long-Range Repulsion Colloidal Model toward Predicting the Viscosity of Protein Solutions. Molecular Pharmaceutics. 2022 Sep 21;19(11):4233–40.

33. Liu Y, Xi Y. Colloidal systems with a short-range attraction and long-range repulsion: Phase diagrams, structures, and dynamics. Current opinion in colloid & interface science. 2019 Feb 1;39:123–36.

34. Liu J, Yin DC, Guo YZ, Wang XK, Xie SX, Lu QQ, Liu YM. Selecting temperature for protein crystallization screens using the temperature dependence of the second virial coefficient. PloS one. 2011 Mar 30;6(3):e17950.

35. Liu J, Nguyen MD, Andya JD, Shire SJ. Reversible self-association increases the viscosity of a concentrated monoclonal antibody in aqueous solution. Journal of pharmaceutical sciences. 2005 Sep 1;94(9):1928–40.

36. Russel WB. The Huggins coefficient as a means for characterizing suspended particles. Journal of the Chemical Society, Faraday Transactions 2: Molecular and Chemical Physics. 1984;80(1):31–41.

37. Salinas BA, Sathish HA, Bishop SM, Harn N, Carpenter JF, Randolph TW. Understanding and modulating opalescence and viscosity in a monoclonal antibody formulation. Journal of pharmaceutical sciences. 2010 Jan 1;99(1):82–93.

38. Meyer B, Hu B, Ionescu R, Hamm C, Wang N, Mach H, Kirchmeier M, Sweeney J. Opalescence of an IgG1 monoclonal antibody formulation is mediated by ionic strength and excipients. BioPharm International. 2009 Apr 1;22(4).

39. Neergaard MS, Kalonia DS, Parshad H, Nielsen AD, Møller EH, Van De Weert M. Viscosity of high concentration protein formulations of monoclonal antibodies of the IgG1 and IgG4 subclass–Prediction of viscosity through protein–protein interaction measurements. European Journal of Pharmaceutical Sciences. 2013 Jun 14;49(3):400–10.

40. Lilyestrom WG, Yadav S, Shire SJ, Scherer TM. Monoclonal antibody self-association, cluster formation, and rheology at high concentrations. The Journal of Physical Chemistry B. 2013 May 30;117(21):6373–84.

41. Arakawa T, Timasheff SN. Stabilization of protein structure by sugars. Biochemistry. 1982 Dec;21(25):6536–44.

42. Back JF, Oakenfull D, Smith MB. Increased thermal stability of proteins in the presence of sugars and polyols. Biochemistry. 1979 Nov 1;18(23):5191–6.

43. Bam NB, Cleland JL, Yang J, Manning MC, Carpenter JF, Kelley RF, Randolph TW. Tween protects recombinant human growth hormone against agitation-induced damage via hydrophobic interactions. Journal of pharmaceutical sciences. 1998 Dec;87(12):1554–9.

44. Wang W. Protein aggregation and its inhibition in biopharmaceutics. International journal of pharmaceutics. 2005 Jan 31;289(1-2):1–30.

45. Mahler HC, Friess W, Grauschopf U, Kiese S. Protein aggregation: pathways, induction factors and analysis. Journal of pharmaceutical sciences. 2009 Sep 1;98(9):2909–34.

46. Singh SM, Bandi S, Jones DN, Mallela KM. Effect of polysorbate 20 and polysorbate 80 on the higher-order structure of a monoclonal antibody and its Fab and Fc fragments probed using 2D nuclear magnetic resonance spectroscopy. Journal of pharmaceutical sciences. 2017 Dec 1;106(12):3486–98.

47. Lauser KT, Rueter AL, Calabrese MA. Polysorbate identity and quantity dictate the extensional flow properties of protein-excipient solutions. AIChE Journal. 2022 Dec;68(12):e17850.

48. Pindrus MA, Shire SJ, Yadav S, Kalonia DS. The effect of low ionic strength on diffusion and viscosity of monoclonal antibodies. Molecular Pharmaceutics. 2018 Jul 11;15(8):3133–42.

49. Li J, Cheng Y, Chen X, Zheng S. Impact of electroviscous effect on viscosity in developing highly concentrated protein formulations: Lessons from non-protein charged colloids. International Journal of Pharmaceutics: X. 2019 Dec 1;1:100002.

50. Yearley EJ, Godfrin PD, Perevozchikova T, Zhang H, Falus P, Porcar L, Nagao M, Curtis JE, Gawande P, Taing R, Zarraga IE. Observation of small cluster formation in concentrated monoclonal antibody solutions and its implications to solution viscosity. Biophysical journal. 2014 Apr 15;106(8):1763–70.

51. Laage D, Elsaesser T, Hynes JT. Water dynamics in the hydration shells of biomolecules. Chemical Reviews. 2017 Aug 23;117(16):10694–725.

52. Garidel P, Blume A, Wagner M. Prediction of colloidal stability of high concentration protein formulations. Pharmaceutical development and technology. 2015 Apr 3;20(3):367–74.

53. Kannan A, Shieh IC, Hristov P, Fuller GG. In-use interfacial stability of monoclonal antibody formulations diluted in saline iv bags. Journal of Pharmaceutical Sciences. 2021 Apr 1;110(4):1687–92.

54. Deiringer N, Friess W. Proteins on the rack: mechanistic studies on protein particle formation during peristaltic pumping. Journal of Pharmaceutical Sciences. 2022 May 1;111(5):1370–8.

55. Tang EM, Virk SS, Underhill PT. Coupling between long ranged repulsions and short ranged attractions in a colloidal model of zero shear rate viscosity. Journal of Rheology. 2022 May 21;66(3):491–504.

56. Sarangapani PS, Hudson SD, Migler KB, Pathak JA. The limitations of an exclusively colloidal view of protein solution hydrodynamics and rheology. Biophysical journal. 2013 Nov 19;105(10):2418–26.

57. Israelachvili JN. Intermolecular and surface forces. Academic press; 2011 Jul 22.

58. Lewus RA, Levy NE, Lenhoff AM, Sandler SI. A comparative study of monoclonal antibodies. 1. Phase behavior and protein–protein interactions. Biotechnology progress. 2015 Jan;31(1):268–76.

59. Tomar DS, Kumar S, Singh SK, Goswami S, Li L. Molecular basis of high viscosity in concentrated antibody solutions: Strategies for high concentration drug product development. InMAbs 2016 Feb 17 (Vol. 8, No. 2, pp. 216–228). Taylor & Francis.

60. Inoue N, Takai E, Arakawa T, Shiraki K. Specific decrease in solution viscosity of antibodies by arginine for therapeutic formulations. Molecular pharmaceutics. 2014 Jun 2;11(6):1889–96.

61. Hirschman J, Venkataramani D, Murphy MI, Patel SM, Du J, Amin S. Application of thin gap rheometry for high shear rate viscosity measurement in monoclonal antibody formulations. Colloids and Surfaces A: Physicochemical and Engineering Aspects. 2021 Oct 5;626:127018.

62. Begum F, Amin S. Investigating the influence of polysorbate 20/80 and polaxomer P188 on the surface & interfacial properties of bovine serum albumin and lysozyme. Pharmaceutical Research. 2019 Jul;36:1–1.

63. Vadodaria SS, Onyianta AJ, Sun D. High-shear rate rheometry of micro-nanofibrillated cellulose (CMF/CNF) suspensions using rotational rheometer. Cellulose. 2018 Oct;25:5535–52.

64. Allmendinger A, Fischer S, Huwyler J, Mahler HC, Schwarb E, Zarraga IE, Mueller R. Rheological characterization and injection forces of concentrated protein formulations: an alternative predictive model for non-Newtonian solutions. European journal of pharmaceutics and biopharmaceutics. 2014 Jul 1;87(2):318–28.

65. Sharma V, Jaishankar A, Wang YC, McKinley GH. Rheology of globular proteins: apparent yield stress, high shear rate viscosity and interfacial viscoelasticity of bovine serum albumin solutions. Soft matter. 2011;7(11):5150–60.

66. Harding SE, Johnson P. The concentration-dependence of macromolecular parameters. Biochemical Journal. 1985 Nov 1;231(3):543–7.

